# Rapid ocular responses are a robust marker for bottom-up driven auditory salience

**DOI:** 10.1101/498485

**Authors:** Sijia Zhao, Nga Wai Yum, Lucas Benjamin, Elia Benhamou, Shigeto Furukawa, Fred Dick, Malcolm Slaney, Maria Chait

## Abstract

Despite the prevalent use of alerting sounds in alarms and human-machine interface systems, we have only a rudimentary understanding of what determines auditory salience - the automatic attraction of attention by sound - and the brain mechanisms which underlie this process. A major roadblock to understanding has been the lack of a robust, objective means of quantifying sound-driven attentional capture. Here we demonstrate that microsaccade inhibition – an oculomotor response arising within 300ms of sound onset - is a robust marker for the salience of brief sounds. Specifically, we show that a ‘crowd sourced’ (N=911) subjective salience ranking correlated robustly with the superior colliculus (SC) mediated ocular freezing response measured in naïve, passively listening participants (replicated in two groups of N=15 each). More salient sounds evoked earlier microsaccadic inhibition, consistent with a faster orienting response. These results establish that microsaccade-indexed activity within the SC is a practical objective measure for auditory salience.

**Significance statement:** Microsaccades are small, rapid, fixational eye movements, measurable with sensitive eye-tracking equipment. We reveal a novel, robust link between microsaccade dynamics and the subjective salience of brief sounds (salience ranking obtained from a large number of participants in an online experiment): Within 300 ms of sound onset, the eyes of naïve, passively listening participants demonstrate different microsaccade patterns as a function of the sound’s perceptual salience. These results position the Superior Colliculus (the generator of microsaccades) as an important brain area to investigate in the context of a putative multi-modal salience-hub and establish a robust objective means for quantifying auditory salience, critical in the development of sound-based human machine interfaces.

## Introduction

Our perceptual experience of our surroundings is governed by a process of competition for limited resources. This involves an interplay between task-focused attention and bottom-up driven processes which automatically bias perception towards certain aspects of the world, to which our brain, through experience or evolution, has been primed to assign particular significance. Characterizing the stimulus features that contribute to such attentional biasing (“bottom-up driven salience”) and understanding the neural processes which underlie involuntary attentional capture - are topics of intense investigation in systems neuroscience (Itti and Koch, 2001, 2000; Kaya and Elhilali, 2014; Kayser et al., 2005). Specifically as it pertains to the auditory modality, the growing use of sound-based human-machine interface systems in public and professional settings (Burrows, 2018) increases the urgency for understanding auditory salience. However, progress in the field has been hindered by the lack of a robust, objective means of measuring attentional capture by sound.

Research in vision has long capitalized on the fact that attentional allocation can be ‘objectively’ decoded from ocular dynamics: Naїve observers free-viewing complex visual scenes tend to demonstrate consistent fixation, saccade and microsaccade patterns that can be used to infer the attributes that attract bottom-up visual attention (Hafed et al., 2009; Krauzlis et al., 2018; Parkhurst et al., 2002; Peters et al., 2005; Veale et al., 2017; Yuval-Greenberg et al., 2014). The underlying network for saccade (and microsaccade) generation is centered on the superior colliculus (SC; Hafed et al., 2009; Krauzlis et al., 2018) with a contribution from the frontal eye fields (FEF; Peel et al., 2016; Veale et al., 2017), consistent with a well-established role for these areas in computing the visual salience map and controlling overt attention (Hafed et al., 2009; Veale et al., 2017; White et al., 2017b, 2017a). The appearance of new events is also associated with a stereotypical ‘ocular freezing’ (also ‘microsaccadic inhibition’; MSI) response—a rapid transient decrease in the incidence of microsaccades, hypothesized to arise through suppression of ongoing activity in the SC by new sensory inputs (Engbert and Kliegl, 2003; Hafed and Clark, 2002; Hafed and Ignashchenkova, 2013; Rolfs et al., 2008). MSI has been shown to systematically vary with visual salience (Bonneh et al., 2014; Rolfs et al., 2008) and recent evidence suggests this inhibition may only occur if observers ultimately become aware of the signal (White and Rolfs, 2016), indicating that the effect reflects the operation of an interrupt process that abruptly halts ongoing activities so as to accelerate an attentional shift towards a potentially survival-critical event.

Sounds also evoke MSI (Rolfs et al., 2008, 2005) in line with a proposal that it reflect the operation of a modality-general orienting mechanism. In fact, sounds cause faster responses than visual stimuli (Rolfs et al., 2008), consistent with the ‘early warning system’ role of hearing (Murphy et al., 2013). However, because only very simple stimuli have been used, the degree to which sound-evoked ocular responses reflect complex, perceptually relevant acoustic properties, remains unknown.

Here we sought to determine whether MSI is sensitive to the perceptual salience of brief sounds. Using a set of environmental sounds, salience-ranked through crowd-sourcing, we show that the ocular-freezing response is systematically modulated by perceptual salience. The results reveal that activity in the SC is modulated by auditory salience, in line with its role as a multi-sensory hub (Stein and Meredith, 1993), and furthermore establish a direct, objective means with which the saliency of brief sounds can be quantified.

## Results

### (1) Crowed-sourced salience ranking yielded a meaningful and stable subjective salience scale

Eighteen environmental sounds (see Sup. materials), originally used by Cummings et al. (2006), were arbitrarily selected for this study. The stimulus set represents the variety of sounds that may be encountered in an urban acoustic environment: including animal sounds, human non-speech sounds (kiss, sneeze), musical instrumental sounds, impact sounds (golf ball, tennis, impact, coins), and an assortment of mechanical sounds (car alarm, alarm clock, camera shutter, pneumatic drill, lawn mower etc.). All stimuli were length (500 ms) and RMS equated. Example spectrograms are in Figure 1A. We obtained salience ranking from 911 online participants via the Amazon Mechanical Turk (MTurk) platform (see Methods). Sounds were presented in pairs and participants were required to report which one was ‘more salient or attention-grabbing’.

#### Salience rating

Over all responses, a small but robust order-of-presentation bias towards the second sound in a pair was observed (mean probability to choose the first sound = 0.47, mean probability to choose the 2^nd^ sound = 0.53; t = −9.240, p<0.001). However, because the order of presentation was counterbalanced across pairs, this bias did not affect the subjective rating results.

To derive a relative measure of salience for each sound, we counted the proportion of pairs on which each sound was judged as more salient, producing a measure of relative salience ranging between 0 and 1 (Figure 1B). It is striking that a clear scale of subjective salience can be captured across these 18 brief, arbitrarily selected, sounds. Variability was estimated using bootstrap re-sampling (1000 iterations), where on each iteration, salience was computed over a subset of the data (see Methods). The error bars in Figure 1B are one standard deviation from this analysis.

**Figure 1:**
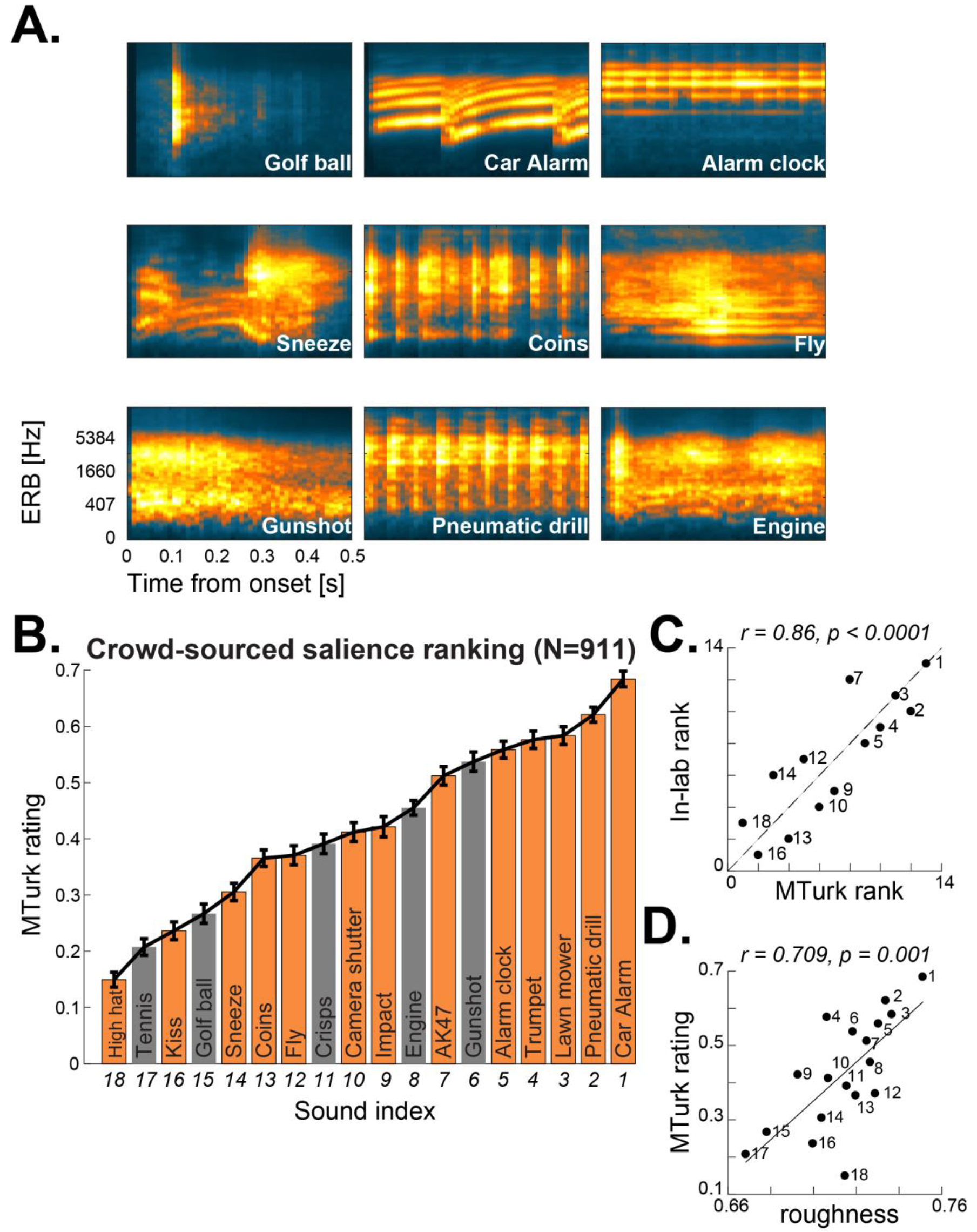
Subjective salience rating for brief environmental sounds. (A) Example spectrograms for a subset of the sounds used in this study (9 out of 18 are displayed). See sup materials for the full dataset (B) Crowdsourced subjective rating collected from MTurk (N=911). The sounds used in the in-lab replication are indicated by orange-coloured bars. Error bars are one standard deviation from bootstrap resampling. (C) Crowdsourced salience rating is strongly correlated with in-lab salience ranking. The dashed line indicates identical ranks (D) Crowdsourced salience rating is strongly correlated with ‘roughness’.

### (2) Crowd-sourced salience scale is strongly correlated with in-lab salience judgements

An in-lab replication was conducted in order to validate the subjective salience scale obtained from the MTurk experiment. The paradigm was essentially identical, but the experiments were performed in a lab setting, and under a controlled listening environment. The main differences were that (1) to reduce test time, sound set was reduced to 13 sounds (indicated in orange bars, Figure 1B; selected to capture the salience range of the full set). (2) All participants listened to all sound pairs during an hour-long session.

The in-lab experiment was analyzed in the same way as described for the MTurk data. The in-lab subjective ranking showed a strong correlation with the online subjective ranking (Spearman r = 0.857, p≤0.0001) (Figure 1C).

### (3) Subjective salience correlates with acoustic roughness

We correlated subjective salience with several key acoustic features, previously hypothesized as contributing to salience:

We found that, despite the fact that **loudness** is known to be a prominent contributor to perceptual salience (Huang and Elhilali, 2017; Kayser et al., 2005; Liao et al., 2016), subjective salience in the present set did not correlate with loudness (p=0.076; see Methods for details about the loudness measure). This may be partly because level was RMS-normalised thus removing some of the larger differences in loudness between sounds.

Next, we tested the relationship between subjective salience and **measures of salience derived from the model of Kayser et al. (2005)**. Several relevant parameters were examined (see Methods). Only correlations with the gradient along the frequency dimension were significant (Spearman’s r=0.525 p=0.027 for the maximum gradient and r=0.488, p=0.049 for the mean gradient; for the rest of the comparisons p>0.155), indicating that perceptual salience may be associated with salience maps in which salient regions are sparsely spread across the spectrum. But this effect did not survive correction for multiple comparisons (across all correlations conducted) and will therefore not be discussed further.

Motivated by previous work (Arnal et al., 2015; Huang and Elhilali, 2017; Sato et al., 2007), we also investigated the correlation between perceptual salience and **roughness** – often quantified by taking the energy in the high end (>30Hz) of the amplitude modulation spectrum (e.g. Arnal et al., 2015). Here, the correlation between crowd-sourced salience and roughness yielded a significant effect (Spearman’s r=0.709, p=0.001; Figure 1D), consistent with accumulating evidence that roughness is a major contributor to salience.

### (4) Crowd-sourced salience correlates with objective measures from ocular dynamics

A subset of 16 out of the 18 original sounds (two sounds, 3 and 9 were excluded due to experimental time constraints) were presented to naїve, centrally fixating, subjects who listened passively to the sounds while their gaze position and pupil diameter were continuously tracked. Sounds were presented in random order, and with a random inter-sound interval between 6 and 7 seconds. Overall, each sound was repeated 20 times across the experimental session. This small number of repetitions was chosen so as to minimize potential effects of perceptual adaptation to the stimuli.

We analyzed two types of rapid orienting responses: the ‘ocular freezing’ (MSI) response (Engbert and Kliegl, 2003; Hafed and Clark, 2002; Hafed and Ignashchenkova, 2013; Rolfs et al., 2008) and the pupil dilation response (PDR; Wang et al., 2017; Wang and Munoz, 2015). Both have been shown to systematically vary with visual salience (Bonneh et al., 2014; Rolfs et al., 2008; Wang et al., 2017, 2014; Wang and Munoz, 2014) and have also been demonstrated to be evoked by auditory stimuli (Liao et al., 2016; Rolfs et al., 2008, 2005; Wang et al., 2014; Wang and Munoz, 2015, 2014; Wetzel et al., 2016), consistent with the notion that they reflect the operation of modality-general interrupt process.

Here we sought to determine whether these responses are modulated by the perceptual salience of brief sounds.

To confirm the robustness and stability of the effects from the first group (group A; N=15), we subsequently repeated the experiment with an additional group of 15 naïve subjects (group B, N=15).

#### Measures of pupillary dilation were not correlated with subjective salience

The temporal evolution of the normalised pupil diameter (the pupil dilation response, PDR) in each participant group is presented in Figure 2. The pupil starts to dilate around 400ms after sound onset, and peaks at approximately 1.16 seconds (ranging from 0.925 to 1.717s). We did not observe significant correlation between subjective salience rating and any key parameters associated with PDR dynamics (see Figure 2 for statistics). This includes (a) the maximum amplitude of the PDR, (b) the PDR peak latency (c) the peak of the PDR derivative (maximum rate of change of the PDR), or (d) PDR derivative peak latency (p≥0.13 for all).

**Figure 2:**
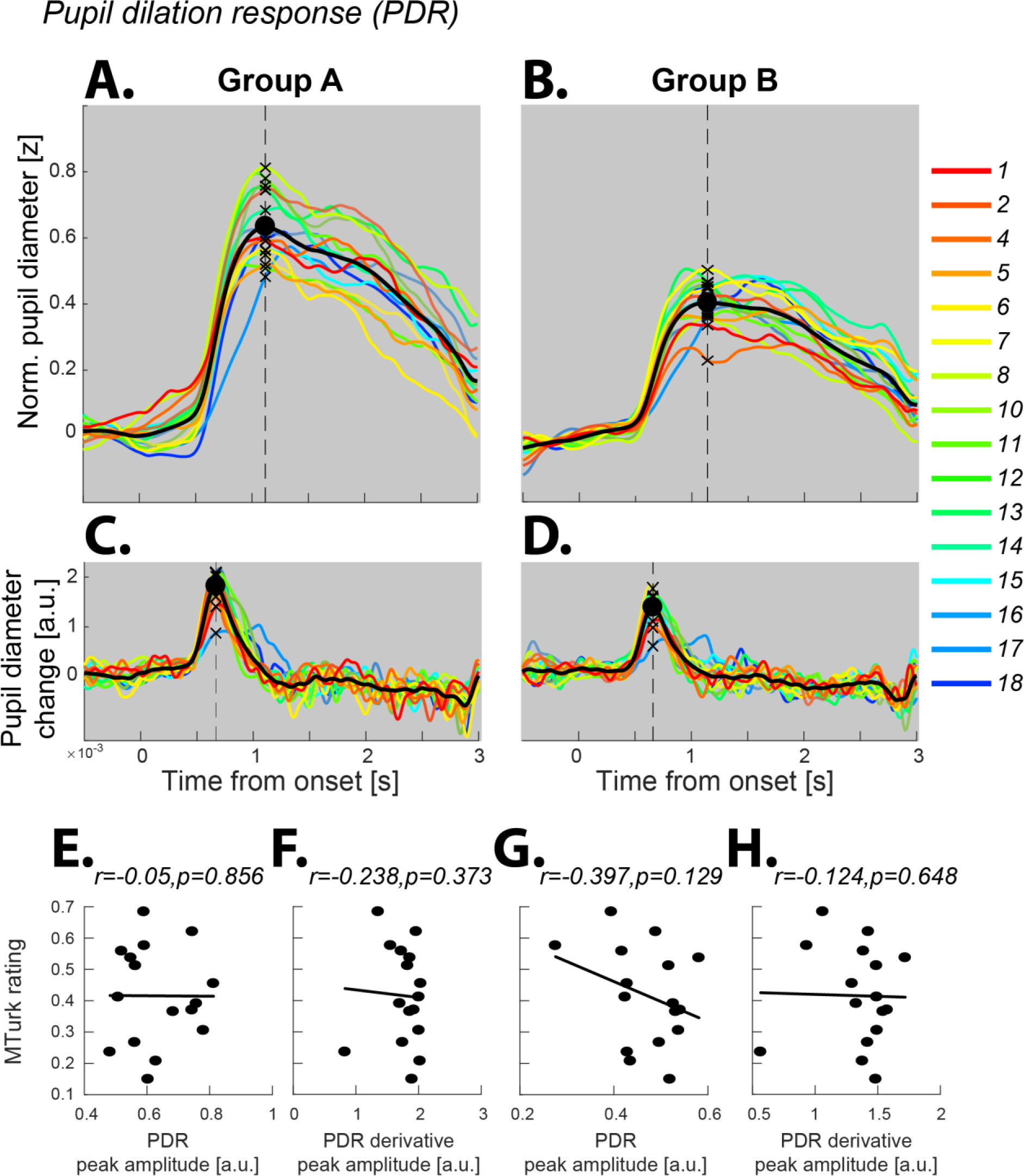
Measures of pupillary dilation are not correlated with subjective salience. Shown are pupil dilation results from (A) Group A and (B) Group B. The solid lines represent the average normalized pupil diameter as a function of time relative to the onset of the sound. The line colour indicates the MTurk salience ranking; more salient sounds are labelled in increasingly warmer colors. The solid black line is the grand-average across all conditions. The dashed line marks the peak average PDR. (C and D) The PDR derivative for both groups. The bottom panel shows the correlations (n.s.) between crowd-sourced salience rating and (E and G) peak PDR amplitude; (F and H) maximum of PDR derivative. None of the effects were significant (including when collapsed across groups; p≥0.822 for all; not shown). A similar analysis based on condition-specific peaks (as opposed to based on the grand average) also did not yield significant effects.

#### Subjective salience is correlated with microsaccadic inhibition

The microsaccade rate obtained from Group A is shown in Figure 3A. In line with previous demonstrations (e.g. Rolfs et al., 2008), we observed an abrupt inhibition of microsaccades following sound presentation. The drop-in rate begins at ~0.3 s after onset and reaches a minimum at 0.45 s. To determine the extent to which these dynamics might vary across the different sounds in the set, we quantified the MSI time for each sound. This was accomplished by computing a grand-mean microsaccade rate time series (averaged across sound conditions; see Figure 3C), identifying its mid-slope amplitude (horizontal dashed line in Figure 3C), and obtaining the time at which the microsaccade pattern associated with each sound condition intersected with this value. This latency (hereafter referred to as the “*microsaccadic inhibition latency*”) strongly correlated with the subjective salience rating (Spearman’s r=−0.704, p=0.002; see Figure S3 for a similar analysis based on the mid-slope time), such that increasing subjective salience was associated with earlier MSI.

To confirm the robustness of this effect, the experiment was repeated with another group of listeners (Group B). The correlation between subjective salience and MSI was replicated (r=−0.579, p=0.021; Figure 3C, bottom). Collapsed across both groups (A+B) we observed r=−0.712 p=0.003. This correlation was significantly different from all PDR correlations reported above (p<0.0001 for all comparisons; see Methods). See Figure S4 for further analyses to confirm effect robustness.

To determine what acoustic information might have driven the observed microsaccade effect, we correlated the MSI latency for each sound (data collapsed across groups), with roughness and loudness estimates computed over the initial 300 ms of each sound (namely, the interval from sound onset and up to the average onset time of ocular inhibition). The correlation with roughness was significant (r=−0.65 p=0.008) and this was also the case for each group separately (Group A: r=−0.7, p=0.007; Group B: r=−0.521, p=0.041). A significant correlation was maintained for shorter intervals down to 150 ms from onset (r=−0.538, p=0.034). This suggests that roughness within the initial portion of the sound might have driven the MS inhibition response. For loudness, only a correlation with the initial 50ms reached significance (r=−0.521 p=0.041) indicating that the loudness at the onset of the snippet may have contributed to the latency of MSI, though this correlation was weak (and not significant for each group separately). A similar analysis based on the measures obtained from the Kayser et al. (2005) model (see Methods) revealed no significant correlations (p≥0.32 for all).

**Figure 3:**
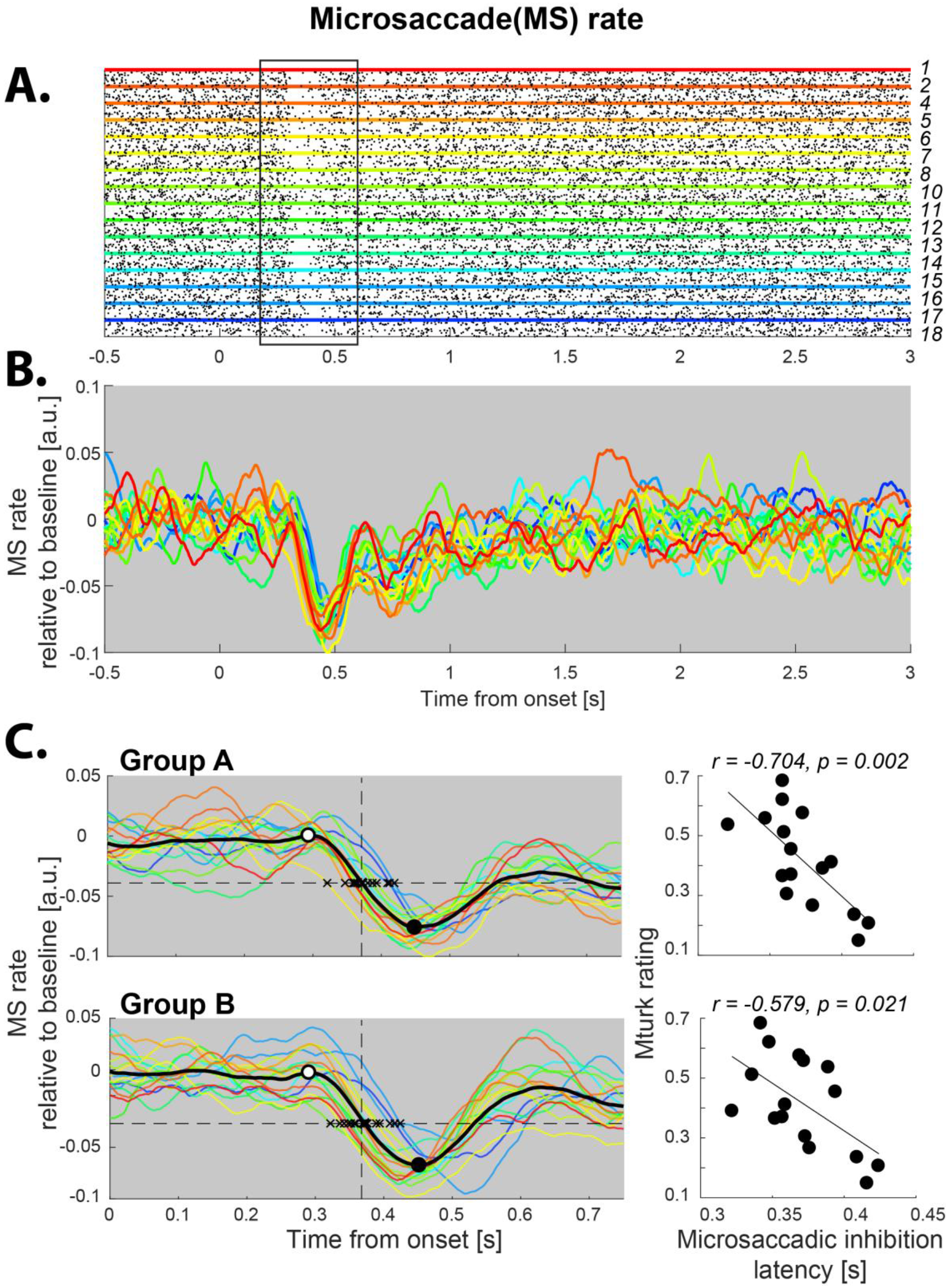
Microsaccadic inhibition (MSI) is correlated with subjective salience. (A) Raster plot of microsaccade events (from group A; pooled across participants) as a function of time relative to sound onset. The Y axis represents single trials; each dot indicates the onset of a microsaccade. Trials are grouped by sound condition and arranged according to the MTurk-derived salience scale (increasingly hot colors indicate rising salience). The region of microsaccadic inhibition, around 0.35-second post-sound onset, is highlighted with a black rectangle. (B) Average Microsaccade rate for each sound (C) focusing on the microsaccade rate during the initial 0.8-second interval after sound onset. The solid black curve is the grand-average microsaccade rate across all conditions. Microsaccadic inhibition commences at approx. 0.3-seconds post sound onset (open circle) and peaks around 0.45 seconds (solid black circle). The horizontal dashed line indicates the mid-slope of the grand average (amplitude = −0.04 a.u., time = 0.372 s, same for both groups). Black crosses mark the time at which the response to each sound intersects with this line, as a measure of MSI latency. On the right are the correlations between these values and the crowd sourced salience rating.

## Discussion

We showed that a ‘crowd sourced’ subjective salience ranking of brief, non-spatial, environmental sounds robustly correlated with the ocular freezing response measured in naїve, passively listening participants. Sounds ranked as more salient evoked earlier microsaccadic inhibition (MSI), consistent with a faster orienting response. These results establish that activity within the superior colliculus (SC) salience hub, arising within ~350 ms after sound onset, is reflective of subjective auditory salience and also demonstrate a practical objective measure for auditory salience.

### Crowd-Sourced Salience

We demonstrated that a robust measure of perceptual salience can be obtained without substantial constraints from a web-based mass-participant experimental platform. Online experimenting is gaining popularity within cognitive science (see Stewart et al., 2017 for a review), including in the auditory modality (Woods et al., 2017; Woods and McDermott, 2018). However, there are various potential drawbacks surrounding this approach, which relate to lack of control over the participants’ listening devices and environment. These may be especially severe for perceptual judgement experiments which demand a high level of engagement from participants. However, the limitations are offset by important unique advantages including the ability to obtain a large amount of data in a short period of time and running brief ‘one-shot’ experiments which are critical for avoiding perceptual adaptation. Furthermore, in the context of salience, the variability of the sound environment may in fact provide ‘real world’ validity to the obtained scale. Here we established that despite the various concerns outlined above, capitalizing on big numbers makes it possible to acquire a stable, informative, salience scale with relatively minimal control of the listeners and their environment. Indeed, the salience scale obtained online correlated robustly with in-lab measures as well as with certain acoustic features previously established as contributing to perceptual salience.

Specifically, we found a strong correlation with ‘roughness’ - the perceptual attribute that is associated with ‘raspy’, ‘buzzing’ or ‘harsh’ sounds. This correlation arose ‘organically’ despite the fact that the sounds in the present study were selected arbitrarily. The link between roughness and subjective salience is in line with previous reports (e.g. Arnal et al., 2015; Huang and Elhilali, 2017; Sato et al., 2007) which established a clear role for this feature in determining the perceptual prominence of sounds. Most recently this was demonstrated in the context of the distinctiveness of screams (Arnal et al., 2015; though Arnal et al. used the term ‘fearful’ in their experiments).

Kayser et al. (2005) have proposed a model for auditory salience, inspired in its architecture by the well-established model for visual salience (Itti and Koch, 2001). We found that none of the parameters derived from that model correlated with the present crowd-sourced scale. This is possibly because the Kayser model is better suited to capturing ‘pop-out’-like saliency, associated with attentional capture by an object which stands out from its background. Instead here we focused on brief sounds reflecting single acoustic sources. We also did not find a correlation with loudness—a well-known contributor to salience (Huang and Elhilali, 2017; Kayser et al., 2005; Liao et al., 2016). The fact that sounds were RMS-equated to control for level likely reduced the perceptual effect of this dimension.

### Acoustic salience does not modulate pupil responses

The pupil dilation response (PDR) indexes activity within the LC-norepinephrine system (Aston-Jones and Cohen, 2005; Joshi et al., 2016), which is proposed to play a key role in controlling global vigilance and arousal (Sara, 2009; Sara and Bouret, 2012). In previous work that reported an association between the PDR and sound salience, the dominant driving feature for the correlation was loudness (Huang and Elhilali, 2017; Liao et al., 2016). In contrast, differences along this dimension were minimized in the present stimuli to allow us to focus on subtler, but potentially behaviourally important, contributors to perceptual salience. Our failure to observe a modulation of the PDR by salience suggests that, at least in the context of auditory inputs, pupil dilation may reflect a non-specific arousal response, evoked by stimuli which cross a certain salience threshold. This account is consistent with the relatively late timing of the PDR (peaking about 1 sec after sound onset) thereby potentially reflecting a later stage of processing than that captured by microsaccades (see below).

### Microsaccadic inhibition (MSI) is a reliable correlate of acoustic salience

We revealed a robust correlation between the latency of microsaccadic inhibition (MSI) and subjective salience: Sounds judged by online raters as more salient were associated with a more rapid inhibition of microsaccadic activity. The effect arose early – from roughly 350 ms after sound onset, pointing to fast underlying circuitry.

Microsaccades are increasingly understood to reflect an active attentional sampling mechanism which is mediated by the SC (Hafed et al., 2015; Krauzlis et al., 2018, 2018; Rolfs, 2009; Rucci and Poletti, 2015; Wang and Munoz, 2015). Accumulating work suggests that their occurrence is not automatic but rather modulated by the general state of the participant and by the availability of computational capacity, such that microsaccade incidence is reduced under high load (Dalmaso et al., 2017; Gao et al., 2015; Widmann et al., 2014; Yablonski et al., 2017). MSI is an extreme case for such an effect of attentional capture on ocular dynamics, interpreted as reflecting an interruption of ongoing attentional sampling so as to prioritize the processing of a potentially important sensory event. The dominant account for MSI is that sensory input to the SC causes an interruption to ongoing activity by disturbing the balance of inhibition and excitation (Rolfs et al., 2008). Previously reported effects of visual salience on microsaccade inhibition (Bonneh et al., 2015; Rolfs et al., 2008; Wang et al., 2017) were therefore interpreted as indicating that visual salience may be coded at the level of the SC (see also Mizzi and Michael, 2014; Veale et al., 2017; White et al., 2017a, 2017b). We showed that the perceptual salience of sounds also modulates this response, consistent with a well-established role for the SC as a multi-sensory hub (Meredith et al., 1987; Meredith and Stein, 1986; Wallace et al., 1998; Wang et al., 2017). Importantly, this effect was observed during a diotic presentation – sounds did not differ spatially and were perceived centrally, within the head.

The brain mechanisms which compute and represent auditory salience remain poorly understood (Kaya and Elhilali, 2014; Kayser et al., 2005). The present results position the SC as a key system to study in this context. There is evidence for projections from the auditory cortex to the SC (Meredith and Clemo, 1989; Zingg et al., 2017) which might mediate the effects observed here, or they may arise via the IC as an intermediary (e.g. Xiong et al., 2015).

Finally, the present experiments focused on the salience of brief sounds presented in silence. However, the ongoing context within which sounds are presented is known to play a critical role in determining their perceptual distinctiveness (Kaya and Elhilali, 2014; Sohoglu and Chait, 2016; Southwell and Chait, 2018). In the future, the paradigm established here can be easily expanded to more complex figure-ground situations or to tracking salience within realistic sound mixtures.

## Methods

### Amazon Mechanical Turk derived subjective salience

The experiment was conducted on the Amazon Mechanical Turk (MTurk) platform - an
online marketplace that allows remote participants (‘workers’ per MTurk nomenclature) to perform tasks via a web-interface. Raters were selected from a large pool of prequalified ‘workers’ in the US who were judged to be reliable over the course of many previous (non-salience related) experiments. Each session is delivered through a ‘human intelligence task’ (HIT) page, which contains instructions, and the stimuli for that session. An example of a HIT page used in the present experiment is in the supplementary materials (Figure S1). Since we were interested in the extent to which a relatively free listening environment can result in meaningful data, we did not impose constraints on sound delivery (e.g. Woods et al., 2017) or level (though it was suggested that the participants listen over headphones).

Sounds were presented in pairs. In total, a pool of 7038 pairs (153 possible pairs × 2 orders to control for order effects × 23 repetitions) were generated. The pairs were then arranged in 207 different HITs by randomly selecting subsets of 34 different pairs from the above pool. Each HIT also included 6, randomly interspersed, ‘catch trials’. On these trials the two sounds were physically identical. Each HIT therefore contained 40 sound pairs, as well as instructions for the task. Each pair had its own ‘Play’ button, which when pressed, started the presentation of the corresponding sound pair, with a 500 ms silent gap between the two sounds. Participants were instructed to report which sound (‘first’ or ‘second’) was “more salient or noticeable. Which sound would you think is more distracting or catches your attention?” Participants could only listen to each pair once before responding by selecting one of the “first”, “second” or “identical” buttons to progress to the next sound pair. Participants were instructed to choose the “identical” button only if the sounds were physically identical (catch trials). Failure to respond appropriately to the catch trials (or choosing the ‘identical’ response for the non - catch trials) indicated lack of engagement with the experiment and resulted in the data from that session being excluded from analysis (overall about 10% exclusion rate, see below). Participants were offered a base pay, calculated relative to the minimum US wage and based on a 5-minute experiment time. To encourage the participants’ engagement, we paid a small bonus based on correctly labeling the identical sounds and subtracted a small amount for each miss. Each HIT was run by 5 unique workers for an overall number of 1035 sessions. The time limit for task completion was set to 60 minutes, though we expected the experiment to last an average of 3 minutes. Figure S2A plots the actual duration distribution. Most sessions were completed within 3 minutes.

Each participant was free to complete up to a maximum of 9 different HITs. A distribution of #HITs per worker is in Figure S2B. Most (71.4%) completed one HIT only, whilst 52 workers (12.4%) took the experiment the maximum number of times. We did not find any interaction between the number of HITs that a worker took and performance on the catch trials. From the total of 1035 sessions completed, 57 included a single missed catch trial, 11 included two missed catch trials and 11 included more than two missed catch trials. Overall 124 sessions were excluded. The remaining sessions were comprised of 384 unique workers, of which 270 completed only one HIT and the rest completed multiple HITs.

Salience ranking was computed by counting the proportion of pairs on which each sound was judged as more salient. Variability was estimated using bootstrap re-sampling (1000 iterations), where on each iteration one session for each of the 207 unique HITs was randomly selected. The error bars in Figure 1B are one standard deviation from this analysis. The same ranking was obtained after removing sessions whose duration exceeded the 90th percentile (14.09min, N=820 remaining) or the 75th percentile (5.98min, N=683 remaining).

#### Acoustic analysis

The salience data were analyzed to examine possible correlations with various acoustic features previously implicated in perceptual salience.

A **loudness** measure was produced by a model in which the acoustic signal was filtered with a bank of bandpass filters of width 1 ERB (Moore and Glasberg, 1983) with centre frequencies spaced 1/2 ERB from 30 Hz to 16 kHz. The instantaneous power of each filter output was smoothed with a 20ms window and elevated to power 0.3 to approximate specific loudness (Hartmann, 1996). Outputs were then averaged across channels. This model was preceded by a combination of high-pass and low-pass filters to approximate effects of outer and middle ear filtering (Killion, 1978).

Several key **measures of salience derived from the model of Kayser et al. (2005)** were examined including: the maximum values within the saliency map, the mean value, and max/mean gradient across the frequency and/or time dimensions.

**Roughness** was calculated from the modulation power spectrum, computed using the approach described in Elliott and Theunissen (2009; see also Arnal et al., 2015). Roughness was quantified as the ratio between power at modulations >30 Hz (30Hz-100 Hz) and those below 30 Hz.

### In-Lab replication

#### Participants

Eighteen paid participants (15 females, average age 23.8, range 18–31) participated this experiment. All reported normal hearing and no history of neurological disorders. None of the participants were a professional musician or known to possess perfect pitch. Experimental procedures were approved by the research ethics committee of University College London, and written informed consent was obtained from each participant.

#### Procedure

As in the MTurk experiment, we used a pairwise task with identical presentation parameters. However, in the lab every participant was presented with the full set of all possible pairs (78 pairs × 2 possible orders × 2 repetitions) for a total of 312 pairs of sounds. These were presented in a random order in six consecutive blocks. Each block also contained 8, randomly interspersed, catch trials (identical sounds). Participants were allowed a short rest between blocks.

The stimuli were delivered to the participants’ ears by Sennheiser HD558 headphones (Sennheiser, Germany) via a UA-33 sound card (Roland Corporation) at a comfortable listening level, self-adjusted by each participant. Stimulus presentation and response recording were controlled with the Psychtoolbox package (Psychophysics Toolbox Version 3; Brainard, 1997) on MATLAB (The MathWorks, Inc.). Participants were tested in a darkened, acoustically shielded room (IAC triple walled sound attenuating booth). Each session lasted for 1 hour, starting with the same instructions given in the MTurk experiment. Participants were instructed to fixate their gaze on a white cross at the centre of the computer screen while listening to the stimuli, and to respond by pressing one of 3 keyboard buttons to indicate ‘sound A more salient’/ ‘sound B more salient’/ ‘identical sounds’. The participant’s response initiated the following trial with a random inter-trial interval of 1.5 to 2 s. Blocks featuring incorrect responses – whether a miss or a false alarm – to any of the eight catch trials indicated lack of engagement and were discarded from analysis; In this instance all participants performed perfectly with a 100% hit rate and 0% false alarm rate resulting in no exclusions.

### Eye tracking

#### Participants

This experiment was performed twice; a total of 30 unique paid participants attended, with 15 participants initially (group A; 14 females; aged 21~28, average 23.53) and a new group of 15 participants subsequently (group B; 14 females; aged 18~30, average 23.13), to replicate the results of the first cohort. No participants were excluded in this experiment. All reported normal hearing and no history of neurological disorders. None were professional musicians or known to possess perfect pitch. All participants were naїve to the aims of the experiment and none had participated in the ‘in-lab’ experiment, above. Experimental procedures were approved by the research ethics committee of University College London, and written informed consent was obtained from each participant.

#### Stimuli and Procedure

Sixteen sounds out of the original set were used in this experiment. Sound #3 and sound #9 were excluded due to experiment length constraints. Participants listened passively to the sounds, presented in random order, and with a random inter sound interval between 6 and 7 seconds. In total, 320 trials (16 sounds × 20 repetitions of each) were presented. Stimuli were diotically delivered to the participants’ ears by Sennheiser HD558 headphones (Sennheiser, Germany) via a Creative Sound Blaster X-Fi sound card (Creative Technology, Ltd.) at a comfortable listening level, self-adjusted by each participant. Stimulus presentation and response recording were controlled with the Psychtoolbox package (Psychophysics Toolbox Version 3; Brainard, 1997) on MATLAB (The MathWorks, Inc.). Participants sat, with their head fixed on a chinrest, in front of a monitor at a viewing distance of 65cm in a dimly lit and acoustically shielded room (IAC triple walled sound-attenuating booth). They were instructed to continuously fixate at a black cross presented at the centre of the screen against a grey background and passively listen to the sounds (no task was performed). A 24 inch monitor (BENQ XL2420T) with resolution of 1920×1080 pixels and a refresh rate of 60 Hz presented the fixation cross and feedback. The visual display remained the same throughout. To avoid confounding effects from the pupillary light reflex, the luminance of the screen and the ambient illuminance of the experimental room were kept constant throughout the experiment. To reduce fatigue, the experiment was divided into nine 4-minute blocks and equivalent-length breaks.

#### Pupil measurement

An infrared eye-tracking camera (Eyelink 1000 Desktop Mount, SR Research Ltd.), positioned just below the monitor, continuously tracked gaze position and recorded pupil diameters, focusing binocularly at a sampling rate of 1000 Hz. The standard five-point calibration procedure for the Eyelink system was conducted prior to each experimental block. Participants were instructed to blink naturally. They were also encouraged to rest their eyes briefly during inter-trial intervals. Prior to each trial, the eye-tracker automatically checked that the participants’ eyes were open and fixated appropriately; trials would not start unless this was confirmed.

#### Across-trial average pupil diameter response

To measure the pupil dilation responses evoked by the sounds, the pupil diameter data of each trial were epoched between 0.5 seconds before to 3 seconds after sound onset. To compare the results across blocks, conditions and participants, the epoched data within each block were normalized to z-scores. A baseline correction was then applied by subtracting the median pupil size over the pre-onset period; subsequently, the data were smoothed with a 150ms Hanning window. Intervals where the eye tracker detected full or partial eye closure were automatically treated as missing data and recovered with shape-preserving piecewise cubic interpolation; epochs with more than 50% missing data were excluded from analysis. On average, less than two trials per participant were rejected.

#### Microsaccade rate analysis

Microsaccade detection was based on the algorithm proposed by Engbert and Kliegl (2003). In short, microsaccades were extracted from the continuous horizontal eye-movement data based on the following criteria: (a) a velocity threshold of λ = 6 times the median-based standard deviation within each block (b) above-threshold velocity lasting for longer than 5ms but less than 100ms (c) the events are binocular (detected in both eyes) with onset disparity less than 10ms (d) The interval between successive microsaccades is longer than 50 ms.

To compute the incidence rate of microsaccade events, the extracted events were represented as unit pulses (Dirac delta). For each sound-condition, in each participant, the event time series were summed and normalized by the number of trials and the sampling rate. Then, a causal smoothing kernel ω(τ) = *α*^2^ × *τ* × *e*^−ατ^ was applied with a decay parameter of 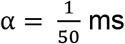 (Dayan and Abbott, 2001; Rolfs et al., 2008; Widmann et al., 2014), paralleling a similar technique for computing neural firing rates from neuronal spike trains (Dayan and Abbott, 2001; see also Joshi et al, 2016; Rolfs et al, 2008). The obtained time series was then baseline corrected over the pre-onset interval. Mean microsaccade rate time series, obtained by averaging across participants for each condition, are used for the analyses reported here (Figure 3A).

### Correlation analysis

Correlations were performed using the Spearman’s rank correlation method. To compare the MSI and PDR-based correlations, we collapsed across groups (N=30) and computed Pearson correlations between the subjective salience and the various eye tracking measures discussed in the results (MSI latency, PDR peak amplitude, PDR derivative peak amplitude). Differences in Pearson correlation coefficients were tested using the procedures for comparing overlapping correlations. The test was conducted through an implementation in the R package cocor (Diedenhofen and Musch, 2015).

#### P-curve analysis

We conducted a p-curve like analysis (Simonsohn et al., 2013) for the purpose of determining the robustness of the main MSI-linked effect. The analysis involved computing a distribution of p values as a function of number of subjects (N). Data were collapsed across the two subject groups (N=30). We iteratively (1000 iterations) selected N samples (5/10/15/30; in different blocks) from this pool (with replacement). For each subset, we computed the correlation between MSI time and subjective salience rating. The resulting p-curves, shown in Figure S4, are significantly right-skewed, with p-values clustered around p=0.01 (Fisher’s Method; p<0.0001; further details in the figure)—indicative of a robust effect. A distribution of associated correlation coefficients (over all significant correlations) is presented on the right of each p-curve.

## Acknowledgements

We are grateful to Makoto Yoneya (NTT) for initiating the pupillometry setup at UCL. This work was supported by an EC Horizon 2020 grant and a BBSRC international partnering award to MC.

## Supplemental Material

**Figure S1:**
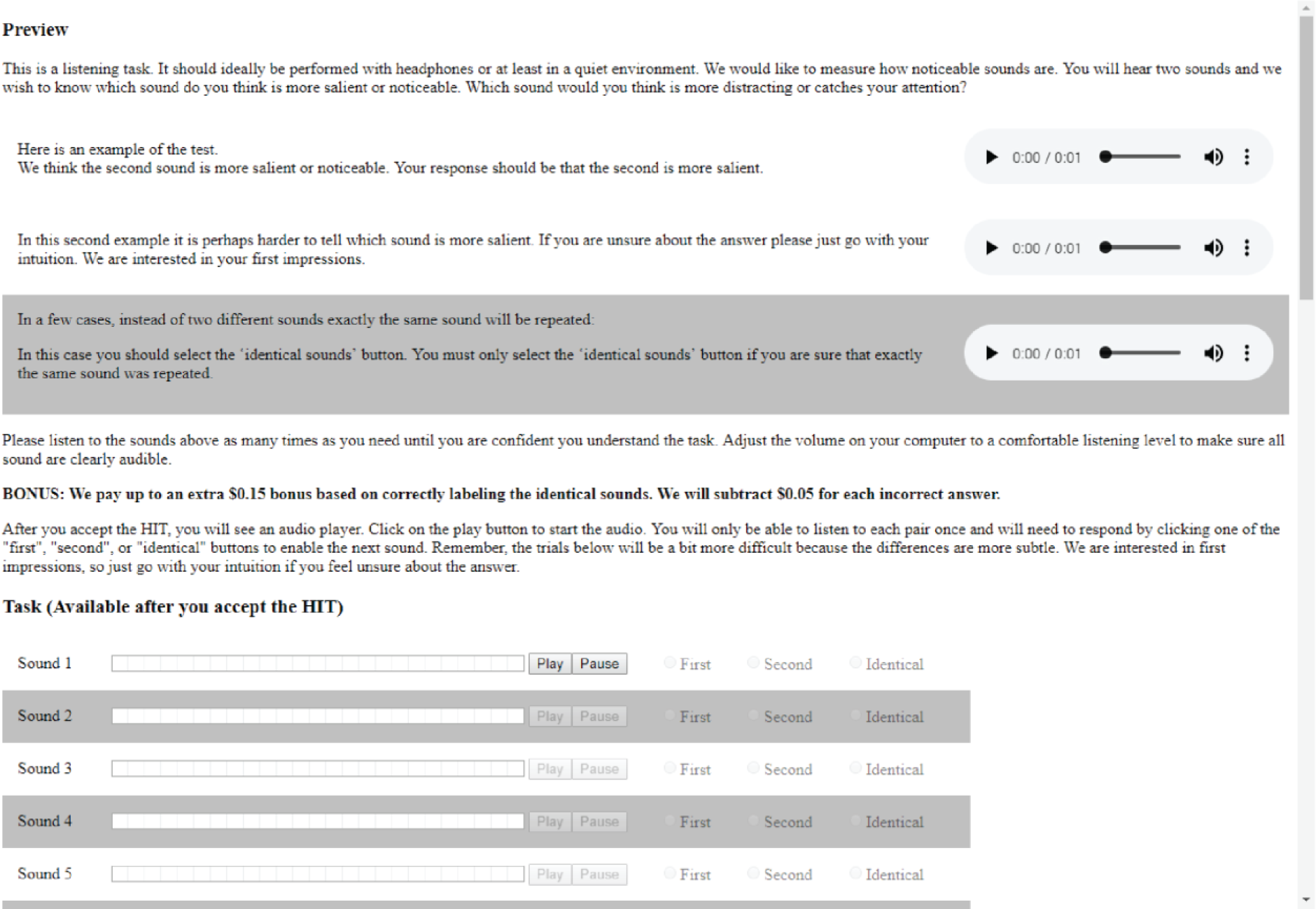
An example of a HIT (‘human intelligence task’) page used in the crowd-sourcing experiment.

**Figure S2:**
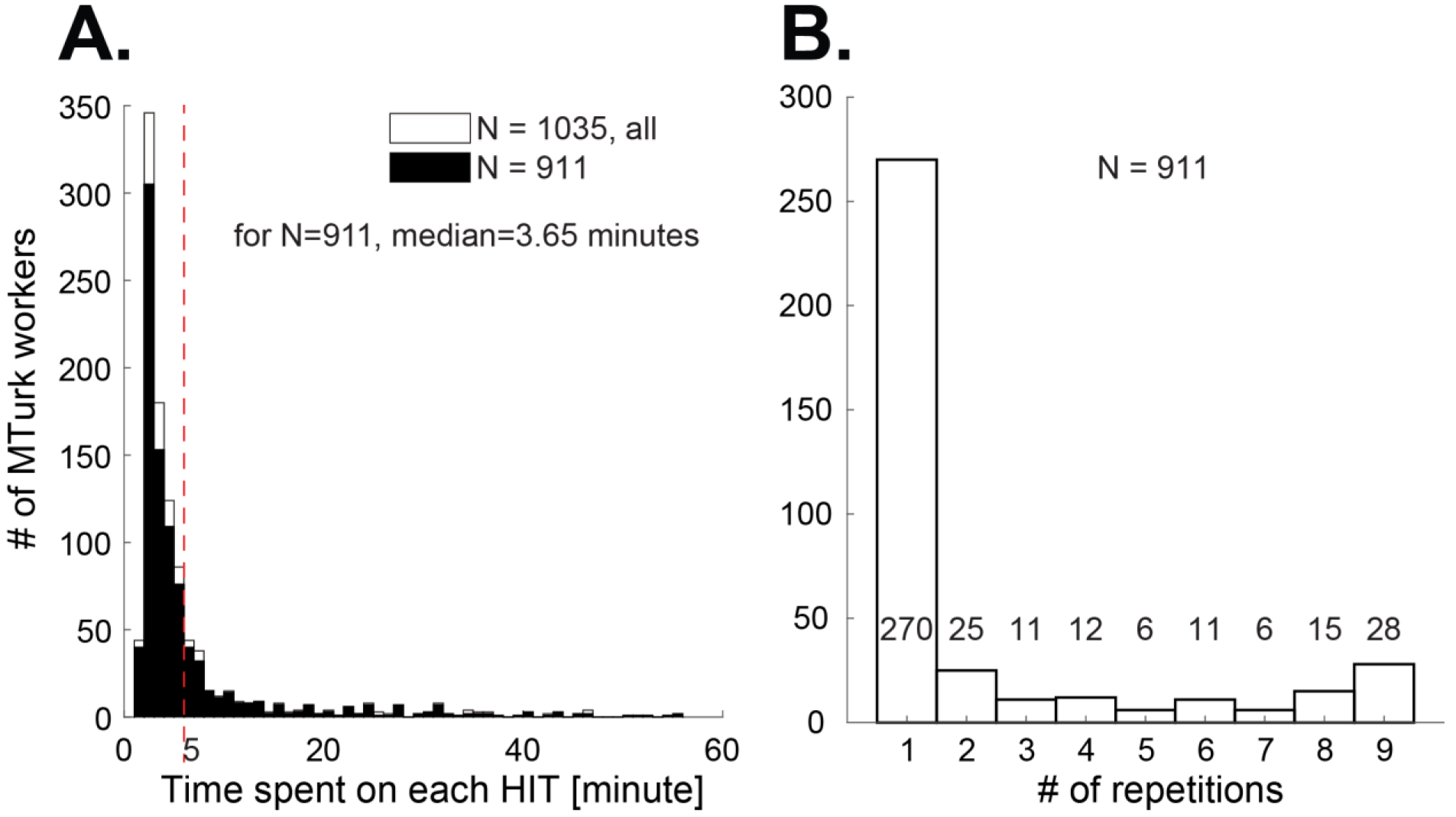
MTurk task data. (A) Distribution of time spent on each HIT. (B) Distribution of number of HITs completed per worker.

**Figure S3:**
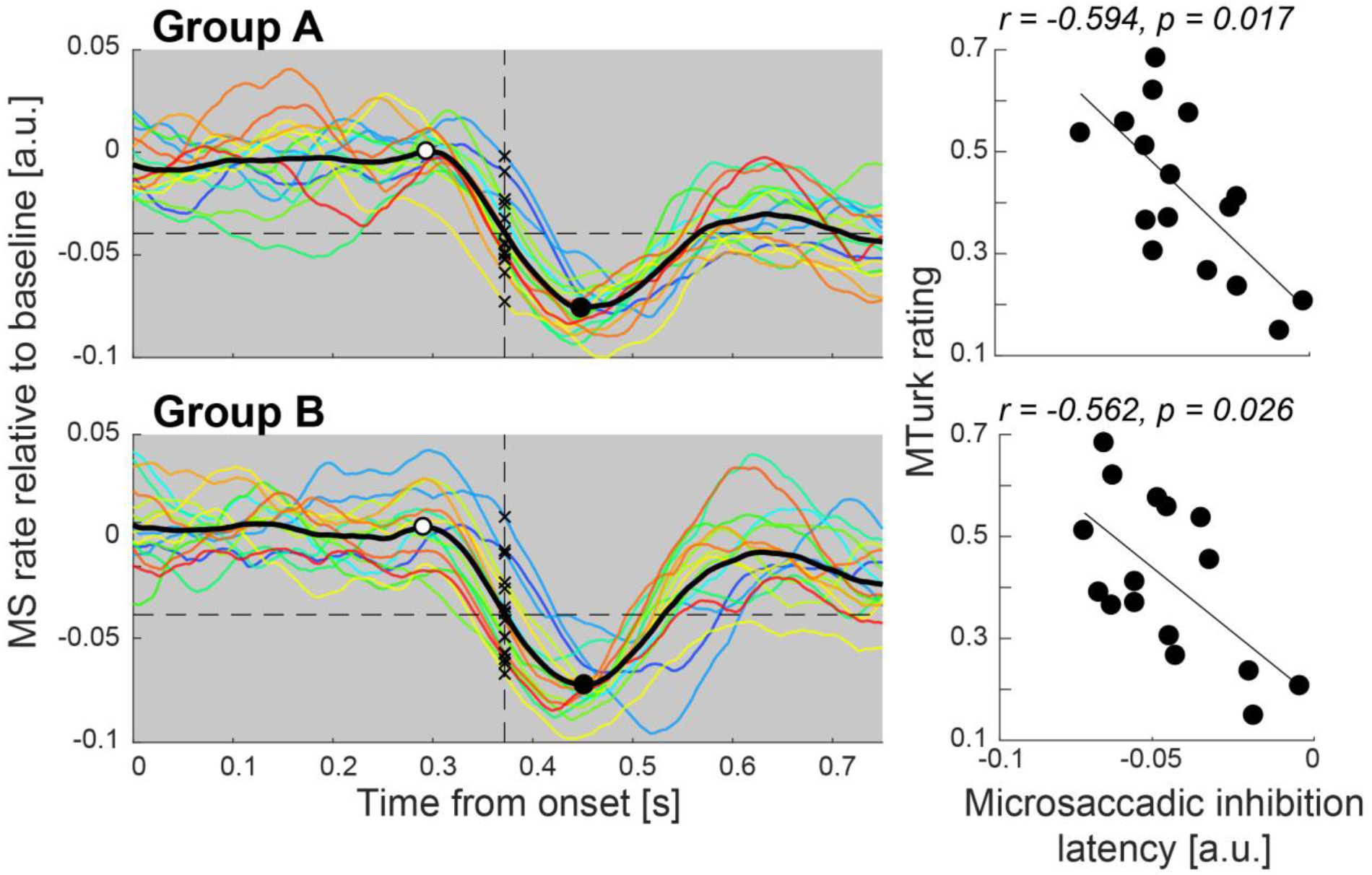
Complementing Figure 3. The dashed lines indicate the mid-slope latency of the grand average. Here black crosses mark the *amplitude* at which the response to each sound intersects with this line, as a measure of MSI. On the right are the correlations between MSI and MTurk-derived salience rating.

**Figure S4:**
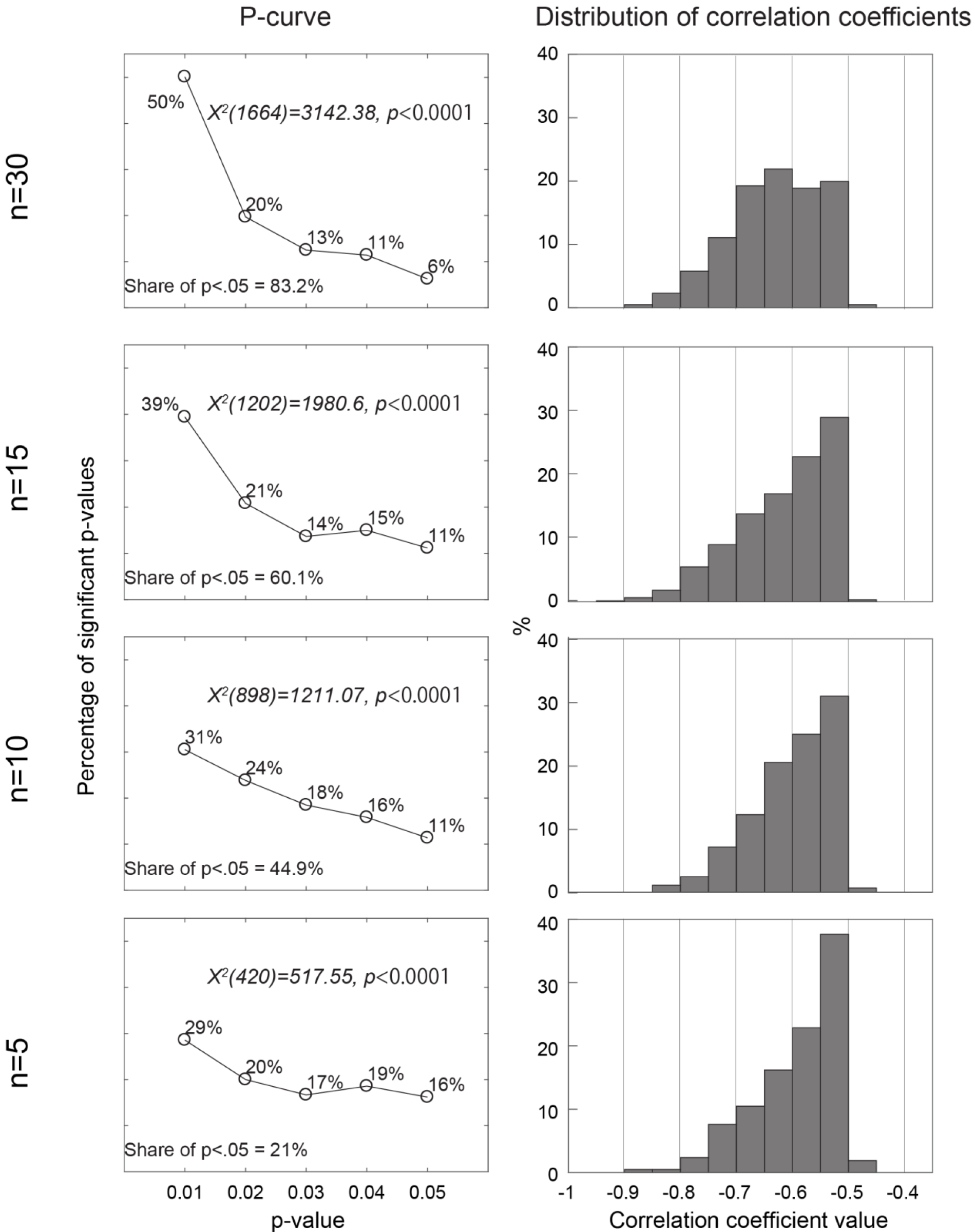
(Left) P-curve analysis for the correlation between MTurk salience rating and MSI as a function of number of participants (N=5; 10; 15; 30). (Right) A distribution of the correlation coefficients associated with each N. Overall, this analysis suggests that to obtain robust effects, =>15 participants are needed. This number of participants offsets the small number of trials (20 trials per sound, randomly intermixed), required to minimize effects of perceptual adaptation to the stimuli.

